# Infection signatures of multiple *Nucleocytoviricota* virus lineages in the brown algae *Undaria pinnatifida* revealed by population-wide genome analysis

**DOI:** 10.1101/2025.02.06.636530

**Authors:** Hiroki Ban

## Abstract

*Undaria pinnatifida* is a large brown algae native to the northwest Pacific. It has been introduced to numerous countries and is listed among the world’s 100 worst invasive species. However, it is also edible and produces various bioactive compounds, making it commercially important. Despite its ecological and economic value, the viruses that infect *U. pinnatifida* remain unknown. In this study, a population-wide genome analysis was performed to investigate viral infection signatures in *U. pinnatifida*. A nearly complete 551-Kb giant endogenous viral element of a *Nucleocytoviricota* virus was identified on a pseudochromosome in the Korean strain genome Upin_Kr2015. This viral region contained viral hallmark genes and auxiliary metabolic genes, including kinases and GDP-mannose dehydrogenase. Additionally, we detected viral hallmark genes in all 43 whole-genome assemblies analyzed, with origins tracing to the orders *Pandoravirales*, *Algavirales*, or *Imitevirales*. Our findings provide strong evidence of frequent *U. pinnatifida* infections by multiple *Nucleocytoviricota* viruses.

## Introduction

The large brown algae *Undaria pinnatifida* (Harvey) Suringar (Laminariales: Phaeophyceae) is a major seaweed native to the northwest Pacific. This species is characterized by its dark-green to brown fronds, which can grow up to 2 m in length. In recent decades, intentional and unintentional dispersal of *U. pinnatifida* have led to its establishment in numerous countries (James et al. 2015; Epstein and Smale 2017), resulting in it being listed among the world’s 100 worst invasive species (Lowe et al. 2000). However, *U. pinnatifida* is also commercially important, accounting for 8% of global annual algal production (FAO 2022). It has been cultivated as an edible seaweed in East Asian countries, including Japan, Korea, and China, for more than half a century (Yamanaka and Akiyama 1993). Additionally, it is recognized in the pharmaceutical and cosmetics industries for its high nutrient content and diverse bioactive compounds (Wang et al. 2018).

Despite its commercial and ecological importance, little is known about viruses infecting *U. pinnatifida*. In general, viral infections in aquaculture cause economic losses by increasing mortality, reducing growth and market value, raising management costs, and affecting ecosystem services (Lafferty et al. 2015). Furthermore, viruses in invasive species can reduce host adaptability to new environments while also potentially infecting native species, leading to serious ecological impacts (Rúa et al. 2011). To better understand the role of viruses in aquaculture and invasion biology, it is essential to elucidate virus–host interactions in *U. pinnatifida*.

The leading candidates for *U. pinnatifida* viruses belong to the phylum *Nucleocytoviricota* (nucleocytoplasmic large DNA viruses; NCLDVs). *Nucleocytoviricota* viruses are double-stranded DNA (dsDNA) viruses with large genomes of 70 Kb to 2.5 Mb (Aylward et al. 2021). These viruses infect a wide range of eukaryotic hosts, from protists to animals, including brown algae (Schulz et al. 2022). Among brown algal viruses, those infecting ectocarpoids (order Ectocarpales), especially *Ectocarpus siliculosus* virus 1 (EsV-1), are the most extensively studied. EsV-1 belongs to the family *Phaeovirinae* of the order *Pandoravirales* (Aylward et al. 2021; Gaïa et al. 2023), which is commonly referred to as phaeoviruses. EsV-1 integrates its genome into the host genome during the free-swimming spore or gamete stage, when host cells lack walls (Maier et al. 2002). This integration enables the viral genome to be inherited by daughter cells and future generations (Müller 1991; Bräutigam et al. 1995; Delaroque et al. 1999). Subsequent studies have identified phaeoviruses in *Laminaria digitata* (McKeown et al. 2017), a species in the same order (Laminariales) as *U. pinnatifida*. A prior study using polymerase chain reaction–based methods found indirect evidence of phaeoviruses in *U. pinnatifida* (McKeown et al. 2018). Additionally, large-scale genomics analyses of brown algae have revealed the insertion of *Nucleocytoviricota* genomes (giant endogenous viral elements; GEVEs) (Moniruzzaman et al. 2020) into multiple brown algal genomes, including that of *U. pinnatifida* (Denoeud et al. 2024).

These findings suggest that *U. pinnatifida* is likely a host for *Nucleocytoviricota* viruses. However, the characteristics and diversity of viruses infecting *U. pinnatifida* remain unknown. Notably, *U. pinnatifida* has two chromosome-level genome assemblies, whole-genome sequences for population genomics analysis, and transcriptomic data from multiple tissues (Shan et al. 2020; Graf et al. 2021). In this study, these datasets are used to investigate virus–host interactions. Results show that a nearly complete genome of a *Nucleocytoviricota* virus is endogenized in the chromosome-level assembly of a Korean strain, a finding overlooked in prior studies. Our results also indicate that *U. pinnatifida* hosts at least three distinct *Nucleocytoviricota* lineages. These findings provide detailed insights into *Nucleocytoviricota* viruses infecting *U. pinnatifida*. This study is anticipated to enhance understanding of virus–host interactions in the context of *U. pinnatifida* ecology and aquaculture.

## Material and Methods

### *Undaria pinnatifida* genome assemblies, whole-genome sequences, and transcriptomes

Two *U. pinnatifida* genome assemblies, the Chinese strain (GCA_012845835.1; ASM1284583v1) and the Korean strain (GCA_020975765.1; Upin_Kr2015), both available in the NCBI database under BioProjects PRJNA575605 and PRJNA646283, were analyzed. Additionally, *U. pinnatifida* DNA and RNA sequences were collected from the Sequence Read Archive (SRA) at NCBI under the same BioProjects. Supplementary Tables 1–3 provide general information on all datasets used.

### Detection of the GEVE region

To identify GEVE regions in *U. pinnatifida* genomes, ViralRecall v2.1 (Aylward and Moniruzzaman 2021) was used with its default parameters. The GEVE region identified on pseudochromosome LG22 in the Upin_Kr2015 assembly was validated as follows. First, guanine and cytosine (GC) content was calculated using a 50-Kb window. Next, corresponding pseudochromosome HiC_scaffold_6 from ASM1284583v1 was aligned using minimap2 v2.28 (Li 2018, 2021) with the “-x asm5” option. To confirm the GEVE region’s connectivity to the host genome, raw long-read sequences from the Korean strain (accession no.: SRR12750387) were mapped to the Upin_Kr2015 genome assembly using minimap2 v2.28 with the “-ax map-pb” option. The resulting SAM file was sorted and converted into BAM file using the “sort” command in SAMtools v1.21 (Danecek et al. 2021) with the “-O BAM” option. The Upin_Kr2015 genome assembly FASTA file and the generated BAM file were indexed using the “faidx” and “index” commands, respectively, in SAMtools v1.21. Integrative Genomics Viewer (IGV) v2.18.4 (Robinson et al. 2011) was employed for visualization.

To determine the *Nucleocytoviricota* virus most closely related to the GEVE region, the average amino acids index (AAI) between the GEVE region and the genomes in the Global Ocean Eukaryotic Viral (GOEV) database (Gaïa et al. 2023) was calculated using EzAAI v.1.2.3 (Kim et al. 2021). The GOEV database includes reference genomes and metagenome-assembled genomes for both *Nucleocytoviricota* and *Mirusviricota*.

### Gene annotation of the GEVE region

Protein-coding genes in the GEVE region were predicted using Prodigal v2.6.3 (Hyatt et al. 2010) with the “-p single” option. To identify *Nucleocytoviricota* hallmark genes, including major capsid protein (MCP), B DNA polymerase (DNApolB), DNA-dependent RNA polymerase subunits A and B (RNApolA and RNApolB, respectively), packaging ATPases (pATPase), D5 DNA primase (Primase), transcription factor II-S (TFIIS), and viral late transcription factor 3 (VLTF3), searches were conducted against eight hidden Markov model (HMM) profiles (Gaïa et al. 2023) using the “hmmsearch” command in HMMER v3.3.2 (Finn et al. 2011) with the “-E 1e-05” option. Additional 149 *Nucleocytoviricota* orthologous genes (Guglielmini et al. 2019) were searched against the HMM profiles (Gaïa et al. 2023) using the “hmmsearch” command in HMMER v3.3.2 with the same “-E 1e-05” option. Protein domains were annotated against Pfam-A v.37.0 using the “hmmscan” command in HMMER v3.3.2 with the “--cut_ga” option.

### Transcriptome of the GEVE region

RNA short read sets from eight different tissues/conditions of the Korean strain (Graf et al. 2021) were subjected to adapter trimming and quality control using fastp v0.23.4 (Chen 2023) with options “-3 -W 6 -M 30 -q 20 -u 50 -n 0 -p −l 50 -w 16”. Each quality-controlled short read set was mapped to the reference genome assembly, Upin_Kr2015, using STAR v2.7.11a (Dobin et al. 2013) with the “--outSAMtype BAM SortedByCoordinate” option. The read depth at each position in the assembly was calculated using the “depth” command in SAMtools v1.21. The number of reads mapped to each gene was determined using HTseq v0.11.2 (Anders et al. 2015). Genes’ transcriptional activities were quantified as transcripts per million, with calculations performed exclusively for genes within the GEVE region.

### Individual assemblies of whole-genome sequences

DNA short read sets from 43 *U. pinnatifida* individuals sampled from China, Korea, France, and New Zealand (Shan et al. 2020; Graf et al. 2021) were subjected to adapter trimming and quality control using fastp v0.23.4 with options “-3 -W 6 -M 30 -q 20 -u 50 -n 0 -p -l 50”.

Quality-controlled short reads were assembled using MEGAHIT v1.2.9 (Li et al. 2015) with the “--presets meta-large” option. Protein-coding genes of each assembly were predicted using Prodigal v2.6.3 with the “-p meta” option.

### Hallmark gene detection and taxonomic assignment

The *Nucleocytoviricota* hallmark genes MCP, DNApolB, RNApolA, RNApolB, pATPase, Primase, TFIIS, and VLTF3 were identified by searching against eight HMM profiles, as described earlier. Protein sequences of viruses in the GOEV database were used as references, with their protein-coding sequences predicted using Prodigal v2.6.3 with the “-p single” option. DNApolB, RNApolA, and RNApolB hit the host orthologous genes, so they excluded from the downstream analysis.

Each retained hallmark gene was assigned a taxonomic rank in the GOEV database using a dual BLAST-based last common ancestor approach (Hingamp et al. 2013). Initially, each hallmark gene’s protein sequence was searched against protein sequences in the GOEV database using the “blastp” command in DIAMOND v2.1.9 (Buchfink et al. 2021) with the “-e 1e-05” option. Aligned regions from the best hits were then re-searched against the same database using the “blastp” command in DIAMOND v2.1.9 with the same option. Hits with lower E-values than those from the initial blastp search were retained, and their LCAs were annotated to the initial queries. Protein sequences without hits in the initial blastp search were annotated as “no hit,” whereas sequences with higher taxonomic ranks than order were annotated as “ambiguous” Taxonomic entries labeled as “NA,” “Na,” “ND,” or “incertae_sedis” in the GOEV database were treated as missing data. Supplementary Table 4 lists the revised taxonomy of the GOEV database.

Most analyses in this study were performed using Julia v1.10.4 (Bezanson et al. 2017). Plots were generated using the Julia packages Makie.jl (Danisch and Krumbiegel 2021) and SankeyMATIC (https://github.com/nowthis/sankeymatic). All other programs used in this study are provided in the Methods section.

## Results

### Continuous 551-Kb GEVE region on a pseudochromosome of the Korean strain

Using ViralRecall to detect *Nucleocytoviricota* signals in the two *U. pinnatifida* assemblies, peaks in the ViralRecall rolling score were identified on pseudochromosome LG22 of the Korean strain (Fig. 1a). A peak in GC content on pseudochromosome LG22 coincided with this region (Fig. 1a), indicating a virus-like feature. Alignment with the corresponding pseudochromosome HiC_scaffold_6 of the Chinese strain showed that this region was absent, confirming it as a bona fide viral insertion in the Korean strain genome (Fig. 1b). Mapping of raw long reads to the Korean strain genome showed that the viral region was connected to the host genome, ruling out misassembly (Fig. S1). These findings demonstrate that a 551-Kb continuous region (1,231,551–1,782,955 bp) on pseudochromosome LG22 represents an integrated GEVE in the *U. pinnatifida* genome.

**Fig. 1.**
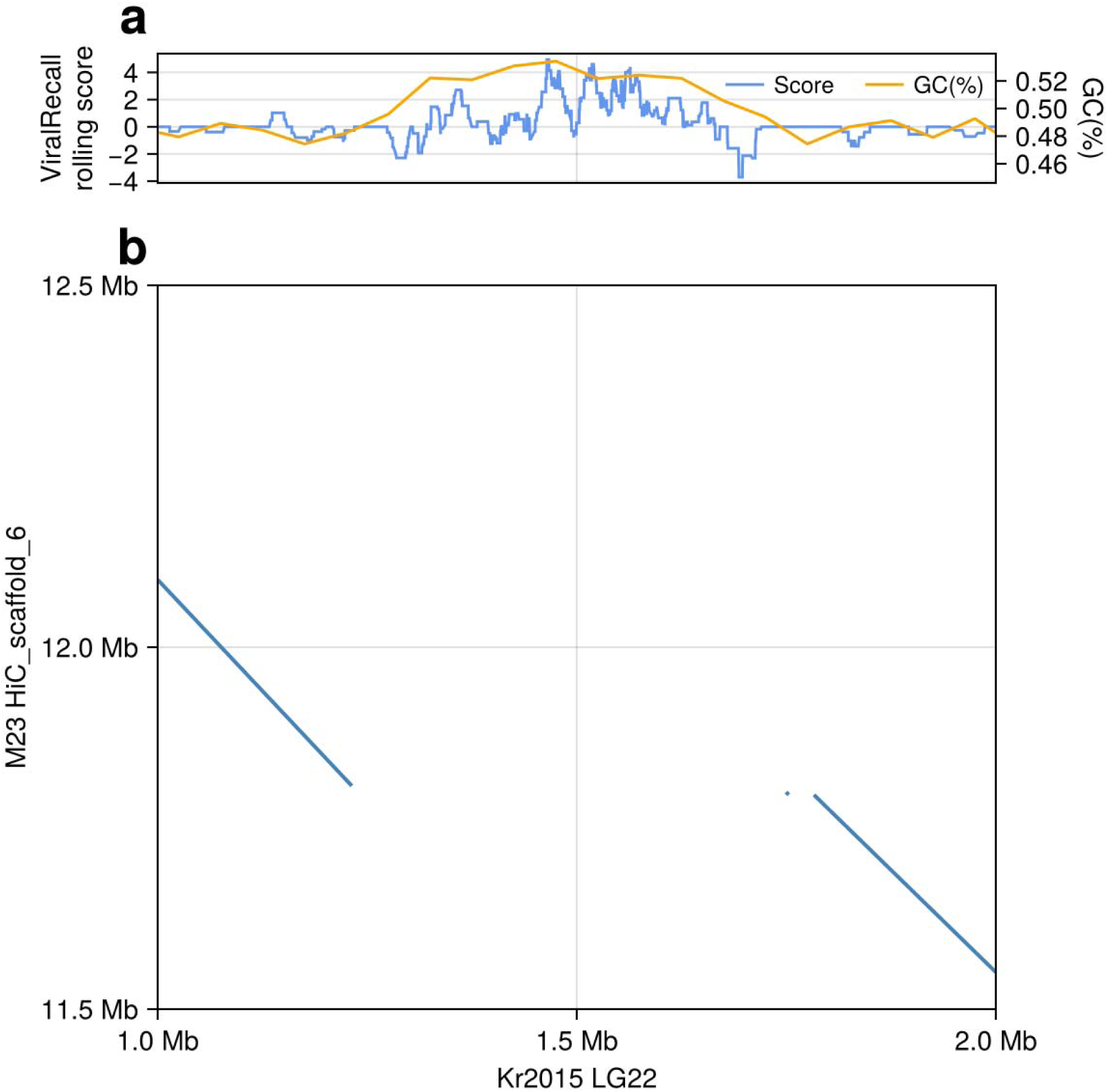
(a) ViralRecall rolling score and GC content along pseudochromosome LG22 of the Korean strain. A positive score indicates a viral signal. GC content was calculated using a 50-Kb window. (b) Dot plot showing the alignment between pseudochromosome LG22 of the Korean strain and the corresponding pseudochromosome HiC_scaffold_6 of the Chinese strain.

### General features of the GEVE

The highest AAI between the GEVE and the genomes in the GOEV database was 67.4%, aligning most closely with EsV-1. The GEVE had a GC content of 51.4%, higher than the 49.8% GC content of pseudochromosome LG22. In total, 651 protein-coding genes were predicted in the GEVE (Fig. 2), with a coding density of 67.8%, lower than the typical >80% found in *Nucleocytoviricota* viruses (Schulz et al. 2020). Among the eight hallmark genes, MCP, Primase, VLTF3, and PolB were clustered in the central GEVE region (Fig. 2), whereas RNApolA, RNApolB, pATPase, and TFIIS were absent. Additionally, 58 *Nucleocytoviricota* orthologous genes were identified (Fig. 2). Among the 97 protein-coding genes functionally annotated using Pfam-A (Fig. 2), such as ribosomal protein S5e with multiple domains (e.g., the tetratricopeptide repeat; Gene ID: 288). The GEVE also contained auxiliary metabolic genes, including those encoding thaumatin family protein, chitin synthase, and GDP-mannose dehydrogenase (Gene IDs: 268, 411, and 412). Additionally, three kinase genes were identified, including one encoding a phytochrome-containing histidine kinase (Gene IDs: 313, 374, and 413). A reverse transcriptase–containing integrase gene was also present (Gene ID: 550).

**Fig. 2.**
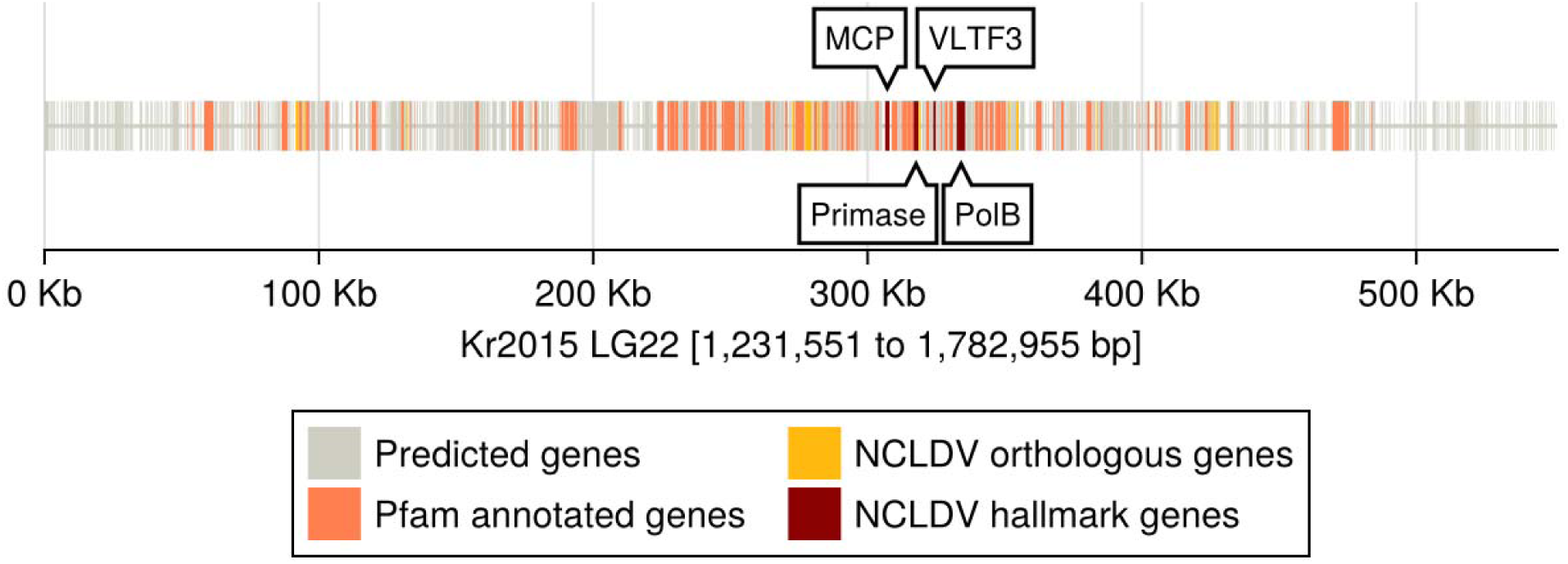
Genomic map depicting the distribution of predicted genes across the 551-Kb region (1,231,551–1,782,955 bp) of pseudochromosome LG22 of the Korean strain. Gray, yellow, orange, and red bars represent predicted genes, *Nucleocytoviricota* (NCLDV) orthologous genes, genes annotated with Pfam-A, and *Nucleocytoviricota* hallmark genes, respectively. Where multiple annotations apply to a gene, priority is given in the following order: hallmark genes, Pfam-A annotations, and *Nucleocytoviricota* orthologous genes. *Nucleocytoviricota* hallmark genes [major capsid protein (MCP), D5 primase (Primase), viral late transcription factor 3 (VLTF3), and B DNA polymerase (PolB)] are marked with arrows indicating their locations.

### Transcriptomes of the GEVE

Mapping of transcriptome reads revealed a distinct expression pattern across all tissues and conditions. The central region of the GEVE was largely silent, whereas both edges were actively transcribed (Fig. 3). This trend persisted when analyzing individual gene expressions (Fig. 4a). Of the 651 genes, 433 showed no expression across all tissues/conditions, whereas 218 were expressed in at least one or more of these tissues/conditions (Fig. 4b). Notably, all *Nucleocytoviricota* hallmark genes and most *Nucleocytoviricota* orthologous genes were not expressed (Fig. 4b). Many of the expressed genes were functionally unknown (Fig. 4b), although some encoded enzymes, such as reverse transcriptase.

**Fig. 3.**
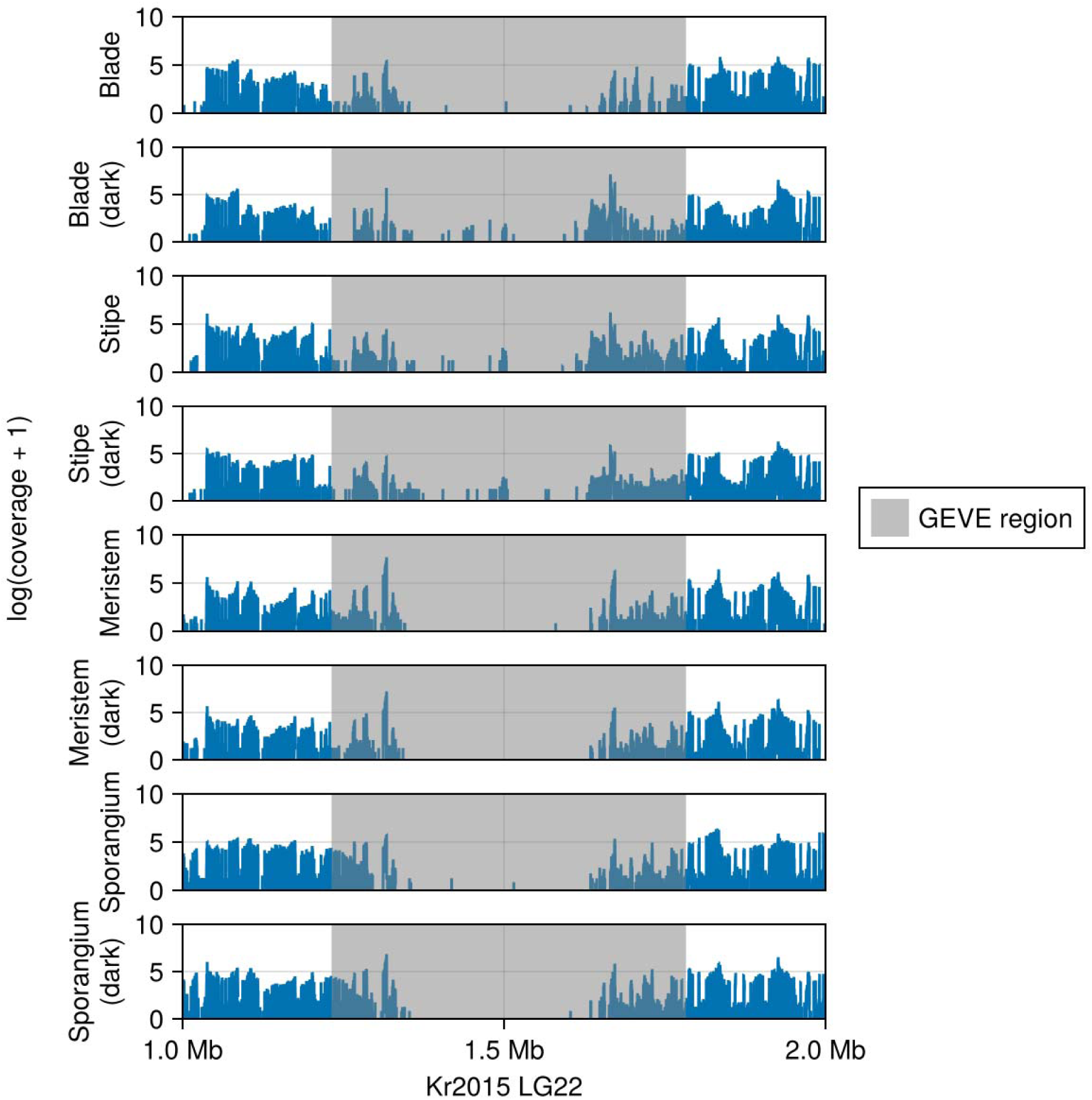
Gene expression across different tissues and conditions in pseudochromosome LG22 of the Korean strain, from 1.0 to 2.0 Mb. Y-axis represent mapped read coverage, transformed to log(‘coverage’ + 1). Panels represent gene expressions in the blade, blade (dark), stipe, stipe (dark), meristem, meristem (dark), sporangium, and sporangium (dark). Gray area highlights the GEVE region (1,231,551–1,782,955 bp).

**Fig. 4.**
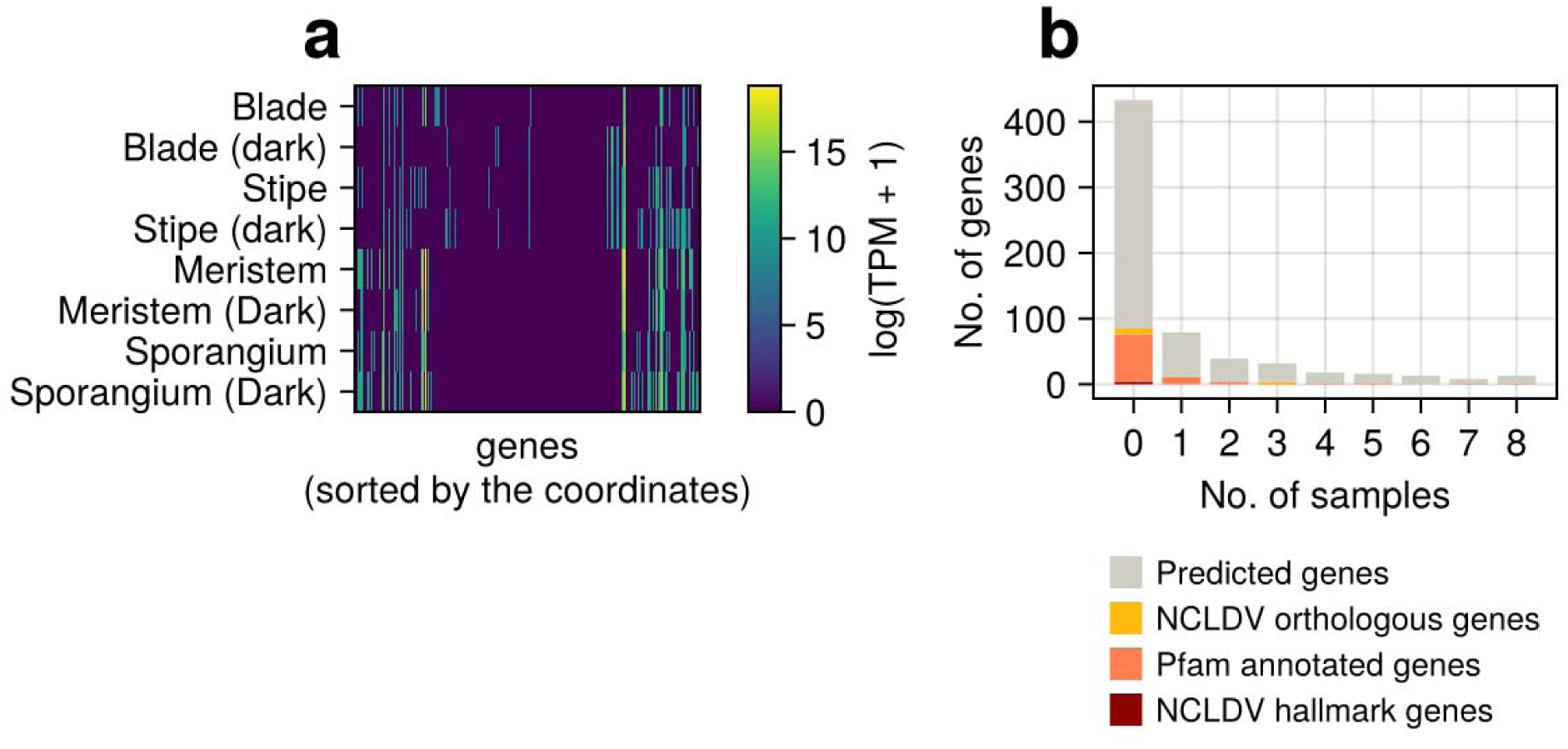
(a) Heatmap representing gene expression across different tissues, with color intensity indicating expression levels from low (blue) to high (yellow). (b) Distribution of detected genes across different samples, categorized by gene type (as shown in Fig. 2). X and Y axes represent the number of tissues/conditions in which the genes are expressed and gene counts, respectively.

### *Nucleocytoviricota* hallmark genes found in all whole-genome assemblies

After assembling each whole-genome sequence individually, 43 assemblies ranging in size from 554 to 890 Kb were obtained. From the predicted protein-coding genes, 282 genes corresponding to 5 hallmark genes (MCP, pATPase, Primase, TFIIS, and VLTF3) were identified. Hallmark genes were present in all assemblies. Specifically, 0–2 MCPs, 1–30 pATPases, 0–11 Primases, 0–2 TFIISs, and 0–5 VLTF3s were identified. MCP was found in 25 assemblies, pATPase in all 43, Primase in 31, TFIIS in 5, and VLTF3 in 27 (Fig. 5). The median protein sequence lengths and scores of these hallmark genes, calculated against the HMM profiles, were lower than those of the viral protein sequences, except for MCP (Fig. 6).

**Fig. 5.**
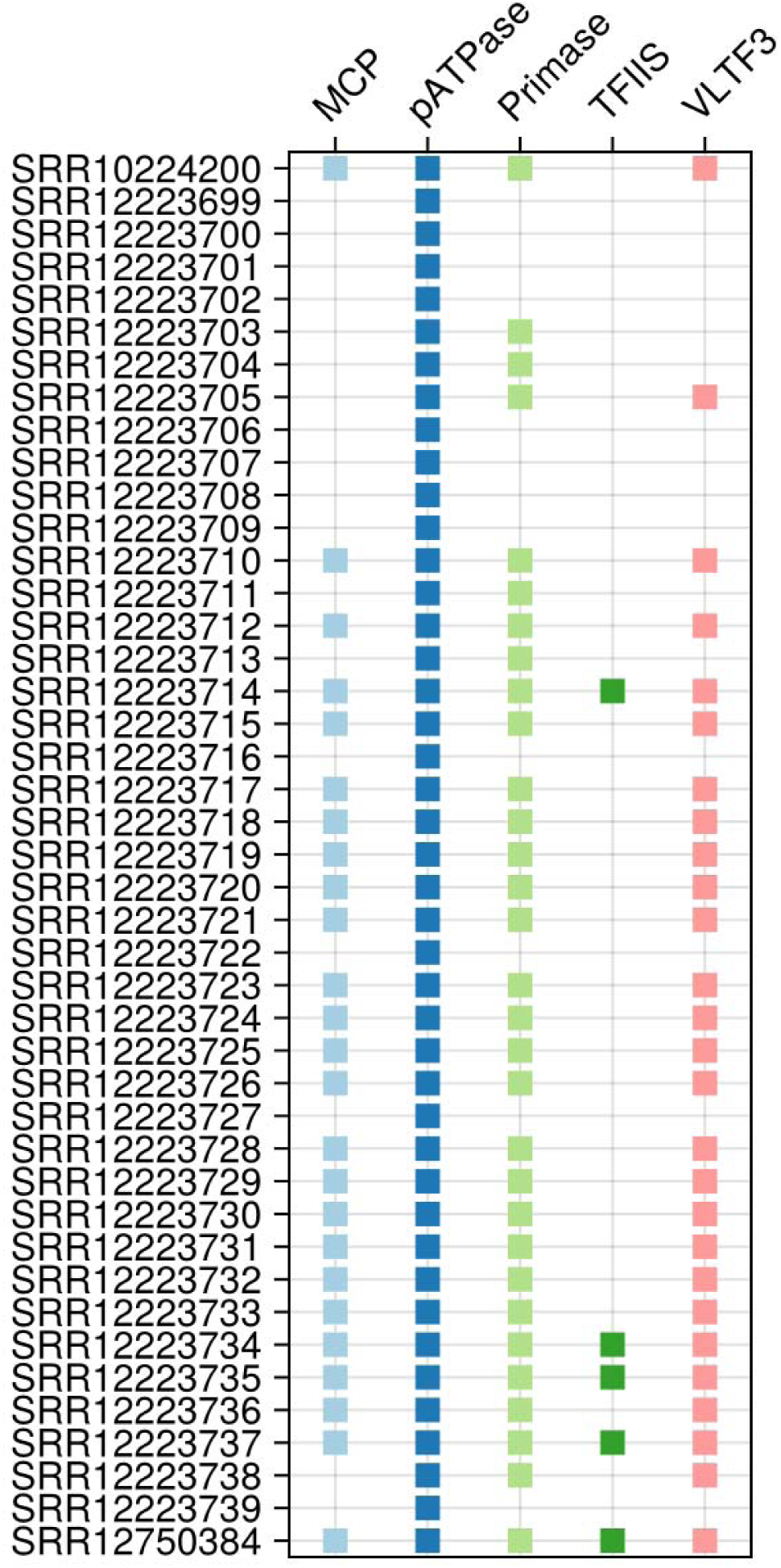
Presence (filled square) or absence of *Nucleocytoviricota* hallmark genes in each assembly. X and Y axes represent *Nucleocytoviricota* hallmark genes and the SRA runs from which the assemblies were derived.

**Fig. 6.**
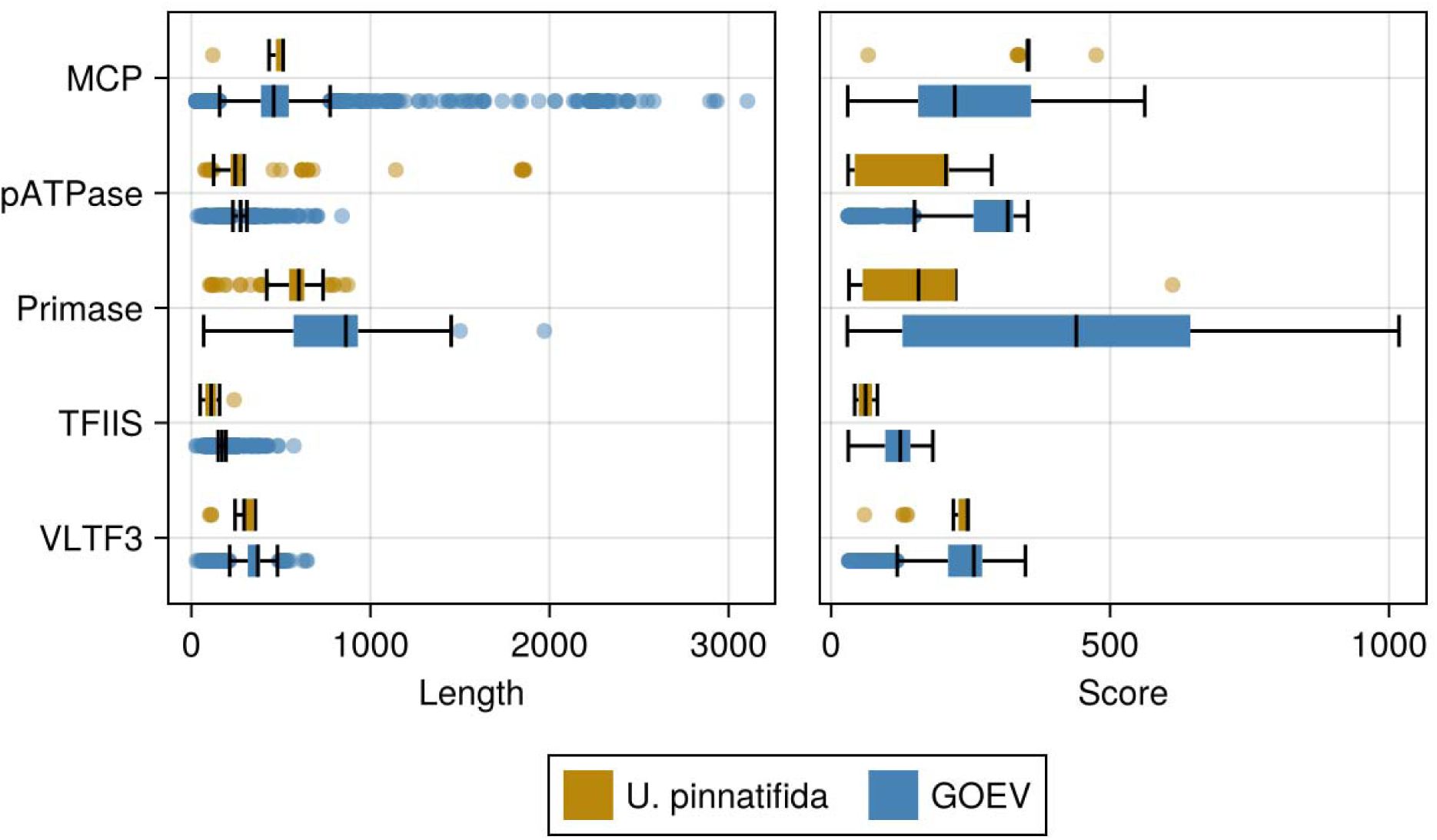
Boxplot showing the sequence lengths and scores of hallmark genes, calculated against HMM profiles, from different sources. *Undaria pinnatifida* assemblies and viral genomes in the GOEV database are shown in brown and blue, respectively.

### Infection signatures of multiple *Nucleocytoviricota* lineages in *U. pinnatifida*

Through taxonomic assignment, 215 of the 282 hallmark genes were classified into specific viral orders within *Nucleocytoviricota* (Fig. 7). No hallmark genes were assigned to *Mirusviricota*, despite its shared hallmark genes with *Nucleocytoviricota*. The majority of genes were assigned to *Pandoravirales*, particularly the family *Phaeovirinae*, with at least one gene from each assembly falling under this classification. In the orders *Algavirales* and *Imitevirales*, hallmark genes were identified in two assemblies each. Within *Algavirales*, a gene from the SRR10224200 assembly was classified under the family “alga_2”, subfamily *Prasinovirinae*, whereas a gene from the SRR12223714 assembly was annotated to the same family but subfamily “alga_2_A.” Within *Imitevirales*, a gene from the SRR12223714 assembly was assigned to the family “basal_clade”, whereas most genes from the SRR12223735 assembly were classified under the family *Mesomimiviridae*, subfamily “meso_4”, genus “meso_4_F2”.

**Fig. 7.**
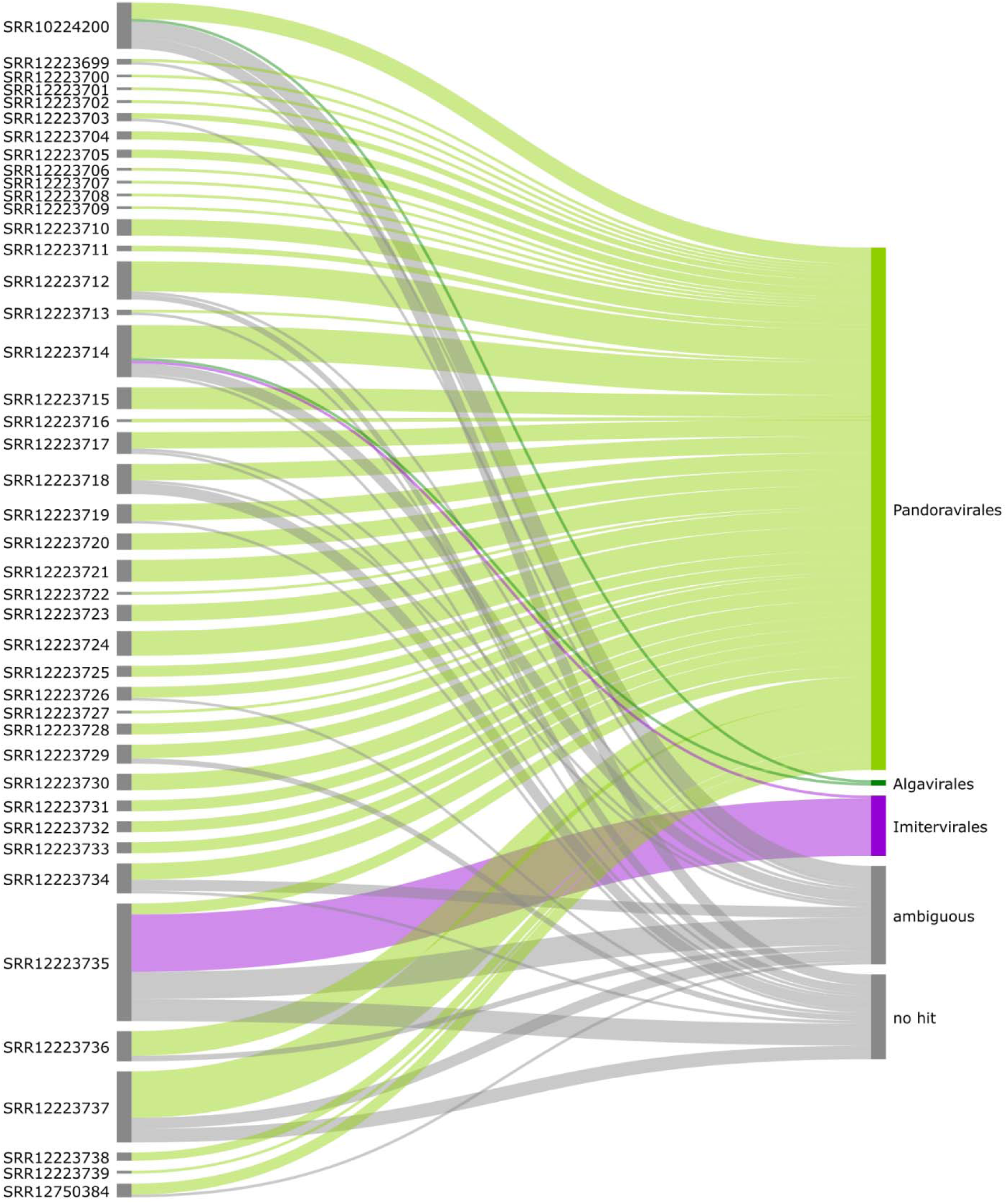
Sankey diagram illustrating the connection between the assemblies of each SRA run and the annotated taxonomy of identified *Nucleocytoviricota* hallmark genes. The “ambiguous” category indicates unclear taxonomy annotation, whereas the “no hit” category includes cases where there was no hit to protein sequences in the GOEV database.

## Discussion

To assess the potential virus–host interactions between *U. pinnatifida* and its associated *Nucleocytoviricota* virus, viral signatures within the genomes of multiple *U. pinnatifida* strains were analyzed. A continuous 551-Kb *Nucleocytoviricota* GEVE region was identified on a pseudochromosome of the Korean strain (Fig. 1 and 2). A previous study predicted the presence of GEVEs at overlapping locations (Denoeud et al. 2024); however, the present study confirmed that the identified region is both longer and continuous. AAI analysis indicated that this GEVE is closely related to the well-characterized phaeovirus EsV-1, suggesting that it likely originated from a member of the phaeoviruses. Although EsV-1 exhibits a lysogenic life cycle (Müller 1991; Bräutigam et al. 1995; Delaroque et al. 1999), the GEVE identified in *U. pinnatifida* appears to be a remnant of a provirus rather than an active virus, as evidenced by its lower coding density compared with typical *Nucleocytoviricota* viruses. This suggests that the region is undergoing pseudogenization and transposon invasion (Zhao et al. 2023). However, its coding density is still higher than that of other known GEVEs, such as the 1.5-Mb continuous GEVE in the arbuscular mycorrhizal fungus *Rhizophagus irregularis*, which has a coding density of only 36.2% (Zhao et al. 2023). This implies that the GEVE insertion in *U. pinnatifida* is relatively recent.

The predicted protein-coding genes within the GEVE offer valuable insights into the original viral lifestyle. The absence of DNA-dependent RNA polymerase genes suggests that the original virus relied heavily on the host’s transcriptional machinery. Notably, genes encoding thaumatin family proteins, commonly involved in stress responses in land plants (Ruiz-Medrano et al. 1992) and possessing known antifungal properties (Vigers et al. 1991), were found within the GEVE. Additionally, a gene for GDP-mannose dehydrogenase, an enzyme responsible for producing GDP-D-mannuronic acid, a key monomer in alginate biosynthesis (with alginate being a component of brown algae cell walls) (Deniaud-Bouët et al. 2017), was also present. These genes and proteins expected to enhance the host’s protection mechanism as a viral extended phenotype (Dawkins 1982). Interestingly, both GDP-mannose dehydrogenase and chitin synthase genes have been identified in the EsV-1 genome, and multiple kinases are known to be encoded in the EsV-1 and *Feldmannia* sp. virus 158 genomes (Delaroque et al. 2001; Schroeder et al. 2009). Therefore, these genes may be common features of phaeoviruses. The GEVE was also found to encode an integrase gene, which the original virus likely used to integrate into the host genome, similar to other phaeoviruses (Delaroque et al. 2001; Schroeder et al. 2009). However, the GEVE integrase gene is unique as it contains a reverse transcriptase. The interaction between an integrase and a reverse transcriptase is an essential step in the integration of retroviruses, such as human immunodeficiency virus type 1 (Tekeste et al. 2015). Notably, no known dsDNA viruses have been shown to integrate into host genomes using both an integrase and a reverse transcriptase, making this a unique and unexplored feature of the original virus.

Consistent expression patterns indicate that the central region of the GEVE was predominantly silent (Fig. 3). This expression pattern led to the suppression of several genes, including *Nucleocytoviricota* hallmark genes, located within the central region of the GEVE (Fig. 2 and 4b). Two potential explanations can be considered for this pattern: first, the virus may have suppressed the expression of hallmark genes, remaining latent until activation; second, the host could be actively suppressing viral gene expression as a defense mechanism. A similar scenario was observed in the 1.5-Mb GEVE region of *R. irregularis*, where chromatin compaction was linked to host defense mechanisms aimed at suppressing gene expression (Zhao et al. 2023). In contrast, genes at the edges of the GEVE, most of which have unknown functions, were actively transcribed (Fig. 3 and 4). These genes may play a role in host metabolism. Additionally, the active expression of reverse transcriptase suggests that it may trigger the horizontal transfer of mobile genetic elements from the virus to the host.

Population-wide genome analysis revealed that individual *U. pinnatifida* strains exhibit evidence of viral infections. The presence of *Nucleocytoviricota* hallmark genes in all assemblies indicates that infections by *Nucleocytoviricota* viruses are frequent in *U. pinnatifida* (Fig. 5). The lengths and lower scores of hallmark genes, excluding MCP, were generally smaller than those of viral reference genes, indicating a lower level of functional constraints on these genes likely underwent pseudogenization after being incorporated into the host genome. Interestingly, MCP did not follow this trend; however, considering the high diversity of *Nucleocytoviricota* MCPs (Krupovic et al. 2020) and the taxonomic bias of viral genes found in *U. pinnatifida* assemblies (Fig. 7), this is likely a statistical artifact rather than a biological anomaly.

Taxonomical assignment of the *Nucleocytoviricota* hallmark genes revealed three primary viral orders: *Pandoravirales*, *Algavirales*, and *Imitevirales* (Fig. 7). Most hallmark genes were annotated to the order *Pandoravirales*, family *Phaeovirinae*, suggesting that the majority of *Nucleocytoviricota* viruses infecting *U. pinnatifida* belong to the phaeovirus group. *Algavirales* and *Imitevirales* viruses were also found to infect *U. pinnatifida*, with this result supported by multiple assemblies, indicating a high level of reliability. These findings align with previous studies showing that *Imitevirales* viruses infect brown algae, supporting the hypothesis that such viruses commonly infect these organisms (Denoeud et al. 2024). However, to the best of our knowledge, this study is the first to provide evidence that *Algavirales* viruses, typically associated with unicellular phytoplankton, also infect brown algae. The genes assigned to *Imitevirales* and *Algavirales* viruses were classified into distinct families, highlighting the diversity of viruses within these orders that infect *U. pinnatifida*.

Through analysis of a large dataset, the present study establishes *U. pinnatifida* as a host for *Nucleocytoviricota* viruses. The investigation of the near-complete GEVE of the Korean strain provided valuable insights into the original viral lifestyle. Transcriptome read mapping revealed consistent expression patterns, shedding light on the GEVE–host relationship. The reassembly of individual whole-genome sequences further supported the frequent occurrence of *Nucleocytoviricota* virus infections in *U. pinnatifida*, providing insights into the diversity of these viruses. However, due to the fragmentation of contigs caused by the short-read assembly, it was not possible to compare the insertion sites of the virus or the lengths of the GEVEs. We believe that our findings in present study will contribute to a deeper understanding of the physiology and diversity of *Nucleocytoviricota* virus–*U. pinnatifida* relationships, with implications for the study of aquatic ecosystems and aquaculture.

## Acknowledgments

Computational time was provided by the SuperComputer System, Institute for Chemical Research, Kyoto University. I thank Drs. Hiromori Shimabukuro, Yuji Tomaru, and Yuki Hongo for their helpful and constructive comments.

## Authors’ contributions

H. Ban: Conceptualization, performing all bioinformatics analyses, writing the original draft and editing manuscript.

## Data Availability

Sequence data used during the current study are available in the NCBI BioProject under accession numbers PRJNA575605 and PRJNA646283. The databases used in the study included GOEV database (Gaïa et al. 2023) and Pfam-A (http://pfam.xfam.org/). The HMM profiles used for analysis of the *Nucleocytoviricota* hallmark genes and additional 149 *Nucleocytoviricota* orthologous genes were obtained from the previous study (Gaïa et al. 2023). Data supporting the present results are available at https://doi.org//10.6084/m9.figshare.28340207.

