## Supplementary Fig. 1 for "Infection signatures of multiple *Nucleocytoviricota* virus lineages in the brown algae *Undaria pinnatifida* revealed by population-wide genome analysis"

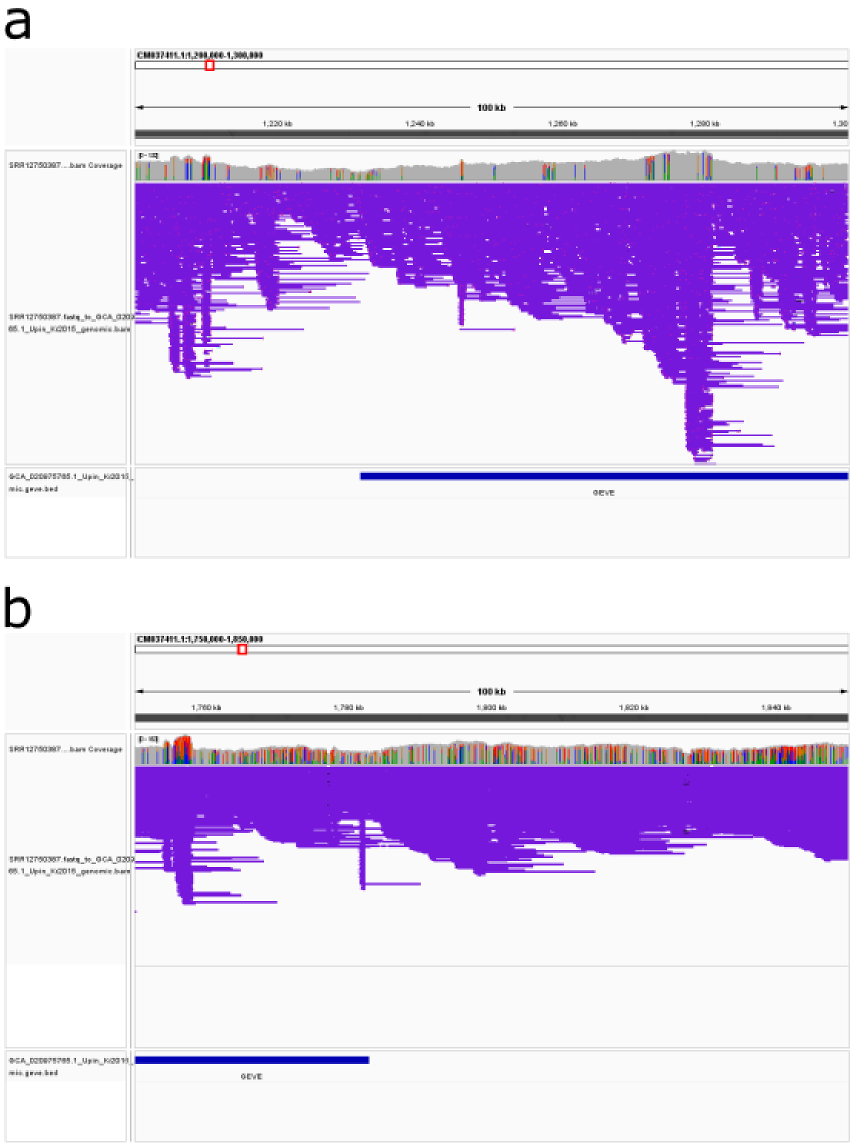


**Supplementary Fig. 1** Screen shot of IGV, representing (a) around 5’ end of the GEVE region (b) around 3’ end of the GEVE region.
